# Signatures of adaptive evolution during human to mink SARS CoV2 cross-species transmission inform estimates of the COVID19 pandemic timing

**DOI:** 10.1101/2021.09.15.459215

**Authors:** Jui-Hung Tai, Shu-Miaw Chaw, Hsiao-Yu Sun, Yi-Cheng Tseng, Guanghao Li, Sui-Yuan Chang, Shiou-Hwei Yeh, Pei-Jer Chen, Hurng-Yi Wang

## Abstract

One of the unique features of SARS-CoV-2 is that it mainly evolved neutrally or under purifying selection during the early pandemic. This contrasts with the preceding epidemics of the closely related SARS-CoV and MERS-CoV, both of which evolved adaptively. It is possible that the SARS-CoV-2 exhibits a unique or adaptive feature which deviates from other coronaviruses. Alternatively, the virus may have been cryptically circulating in humans for a sufficient time to have acquired adaptive changes for efficient transmission before the onset of the current pandemic. In order to test the above scenarios, we analyzed the SARS-CoV-2 sequences from minks (*Neovision vision*) and parenteral human strains. In the early phase of the mink epidemic (April to May 2020), nonsynonymous to synonymous mutation ratios per site within the spike protein was 2.93, indicating a selection process favoring adaptive amino acid changes. In addition, mutations within this protein concentrated within its receptor binding domain and receptor binding motif. Positive selection also left a trace on linked neutral variation. An excess of high frequency derived variants produced by genetic hitchhiking was found during middle (June to July 2020) and early late (August to September 2020) phases of the mink epidemic, but quickly diminished in October and November 2020. Strong positive selection found in SARS-CoV-2 from minks implies that the virus may be not unique in super-adapting to a wide range of new hosts. The mink study suggests that SARS-CoV-2 already went through adaptive evolution in humans, and likely been circulating in humans at least six months before the first case found in Wuhan, China. We also discuss circumstances under which the virus can be well-adapted to its host but fail to induce an outbreak.

## INTRODUCTION

The pandemic coronavirus disease 2019 (COVID-19) which was first recorded in the city of Wuhan, China, is caused by the severe acute respiratory syndrome coronavirus 2 (SARS-CoV-2) [1]. SARS-CoV-2 is the seventh coronavirus found to infect humans. Among the other six, SARS-CoV and MERS-CoV can cause severe respiratory illness, whereas seasonal 229E, HKU1, NL63, and OC43 produce mild symptoms [2]. SARS-CoV-2 exhibits 96% similarity to a coronavirus collected in Yunnan Province, China from a bat, *Rhinolophus affinis*. It is, therefore, possible that the virus may have a zoonotic origin from bats [3, 4].

For a cross-species transmitted virus to achieve high infectiousness in the new host, multiple changes, each of conferring a selective advantage, are necessary [5–7]. For example, a series of small incremental adaptations appear to underly the emergence of SARS-CoV and MERS-CoV that infect humans [8, 9]. Both SARS-CoV and SARS-CoV-2 use their spike protein to mediate entry into host cells. This protein first binds to its host receptor, angiotensin converting enzyme 2 (ACE2), and subsequently mediates the fusion of the viral and host membranes. This receptor binding is the first and one of the most important steps in viral infection of host cells. During the short epidemic in 2002-2003, several rounds of adaptive changes have been documented, especially in the spike protein, in SARS-CoV genomes [10, 11]. Analysis of MERS-CoV sequences also revealed that genetic variability of the spike gene was shaped by multiple adaptive changes [12, 13].

Nevertheless, in the first seven months of the current pandemic (December 2019 - June 2020), SARS-CoV-2 has predominantly evolved under neutral or purifying selection [14–16]. The only probably exception is the D614G mutation in the spike protein that increases viral transmissibility [17, 18]. Although some ORFs such as orf3a and orf8 showed Ka/Ks > 1 in the early pandemic [15], it was due to co-segregation of both ancestral and derived alleles, such as G215V (orf3a) and L84S (orf8), at the same time. As these derived alleles finally went extinct, it is unclear if they were in fact adaptive.

The lack of a positive selection signature in the early SARS-CoV-2 pandemic is in contrast to its precedents, i.e., SARS-CoV and MERS-CoV. It is possible that the SARS-CoV-2 exhibits a unique or preadapted feature distinguishing it from other coronaviruses [19], and enables its efficient cross-transmission to humans and other species without altering its genome. Alternatively, SARS-CoV-2 may have been cryptically circulating in humans for some time before being noticed. During that period, the virus may have acquired adaptive changes to efficiently transmit among humans. After adapting to the new host, most RNA viruses exhibit strong negative selection [20]. Therefore, the signature of positive selection may have become obscured at the time the pandemic took off.

These two scenarios can be tested by examining evolutionary patterns of the virus causing epidemics in other species. If SARS-CoV-2 is preadapted for cross-species transmission, positive selection should not be expected. Otherwise, if SARS-CoV-2 has experienced adaptive evolution to cause the pandemic in humans, signatures of accelerated adaptation should be revealed when it jumps to other species. The transmission of SARS-CoV-2 from humans to minks (*Neovision vision*) thus provides an excellent opportunity to test these scenarios. In this study, we analyzed the sequences from SARS-CoV-2 viruses that infected minks. Our results show a strong signature of positive selection during the early epidemic, with the signal rapidly diminishing later in the outbreak. We also discuss how the virus can have circulated within human populations without being noticed while accumulating adaptive changes.

## MATERIALS and METHODS

### Data Collection

All sequences were downloaded from the Global Initiative on Sharing Avian Influenza Data (GISAID, https://www.gisaid.org/) on or before 2021/2/5. Only complete and high coverage genomes were used. All 796 SARS-CoV-2 genomes labeled as *Neovision vision* (minks) from Denmark, Netherlands, USA, Poland, and Canada were included. We also retrieved all human SARS-CoV-2 sequences from the Netherlands (6625), Poland (406), and Canada (7102). For Denmark and USA, due to the sheer amount of data available, only sequences collected between dates that are seven days before the first mink sequence and seven days after the last mink sequence were included. As a result, 27,971 SARS-CoV-2 genomic sequences were used.

For early-stage data, a collection of 1476 complete and high coverage genomic sequences, with the collection starting from the earliest sequence to February (2019/12/24 ~ 2020/02/29), were retrieved.

For B.1.351 (a.k.a. β or South African strain), we downloaded complete and high coverage sequences that were labeled as lineage B.1.351 of the GH clade and collected before 2020/12/31 on 2021/4/26. The rest of the 1117 non-B.1.351 sequences from South Africa, submitted before 2020/12/31, were also downloaded. For B.1.1.7 (a.k.a. α or English) strain, we downloaded complete and high coverage sequences that were labeled as lineage B.1.1.7 of the GRY clade and collected before 2020/11/30 on 2021/4/26. The rest of the 63492 non-B.1.1.7 sequences from England, submitted before 2020/12/31 and collected between August and November, were also downloaded.

### Sequence analyses and phylogenetic reconstruction

All sequences were aligned against the reference genome (EPI_ISL_402125) using the default settings in ClustalW [21]. Phylogenies were constructed using IQ-TREE 2.1.2 [22]. Number of nonsynonymous changes per nonsynonymous site (Ka) and synonymous changes per synonymous site (Ks) among genomes were estimated based Li-Wu-Luo’s method [23] implemented in MEGA-X [24]. Kimura’s two-parameter model was used to estimate genetic distance between sequences.

For site frequency spectrum (SFS) construction, sequence sets that were immediate sister groups to target groups, including mink-1, B.1.1.7, and B.1.351, based on the phylogenies were used to infer directionality of changes. For the early-stage SARS-CoV-2 sequences, we first used RaTG13 as an outgoup to construct the SFS. We also crossreferenced the directionality of changes based on phylogeny and date of collection. To test whether the observed SFS deviates from neutral expectation under exponential population growth, a custom R script was used based on the Durrett theorem [25].

The ancestor sequences of SARS-CoV-2 and RaTG13 were reconstructed using codeml implemented in PAML 4 [26] under the free ratio model. The sequences used for this analysis included Rf1 (DQ412042.1), HKU3-1 (DQ022305.2), BM48-31 (NC_014470.1), ZC45 (MG772933.1), and ZXC21 (MG772934.1). The reconstructed ancestor sequence was used to infer nucleotide changes after the divergence of SARS-CoV-2 and RaTG13.

### Positive selection in SARS-CoV-2

To examine signatures of positive selection in the SARS-CoV-2 isolates derived from minks, we included all sequences from Netherland. In order to facilitate our analyses, we only retained sequences derived from humans with less than 99.9% nucleotide identify. For SARS-CoV-2 from minks, sequences with ambiguous nucleotides were removed. The resulting dataset contains 92 sequences (32 humans and 60 minks). A maximum likelihood tree was constructed using MEGA-X.

An array of selection detection methods implemented in HyPhy was applied to detect whether the lineage leading to minks has experienced adaptive evolution [27]. W used the fixed effects likelihood (FEL) method [28] to infer amino acid sites under positive selection within minks. We also searched for evidence of positive selection on specific branches using the adaptive branch-site random effects likelihood (aBSREL) method [29]. Because identical or essentially identical sequences do not increase power for codon-based methods to detect selection, we set genetic distance of 0.0005 for B.1.351 and 0.001 for B.1.1.7 for the above analyses.

## RESULTS

### Adaptive evolution of SARS-CoV-2 from Minks

The phylogeny of SARS-CoV-2 derived from minks and their parenteral human strains is shown in (Fig. 1a) (Materials and Methods). Human to mink transmission clearly occurred multiple times, the majority of these events failing to trigger an epidemic. Because selection has been continuously operating in the human hosts, it is reasonable to expect higher infectiousness in humans than in minks [5]. Among all inter-species transmission events, we observed three clusters of infections (mink-1 to 3, all from the Netherlands), suggesting the emergence of new SARS-CoV-2 strains that can efficiently infect minks (colored clade in Fig. 1a). One of the clusters (mink-1) lasted for more than six months (Fig. 1b), implying that the strain may have acquired new mutations to sustain its infection in the new host.

**Fig. 1.**
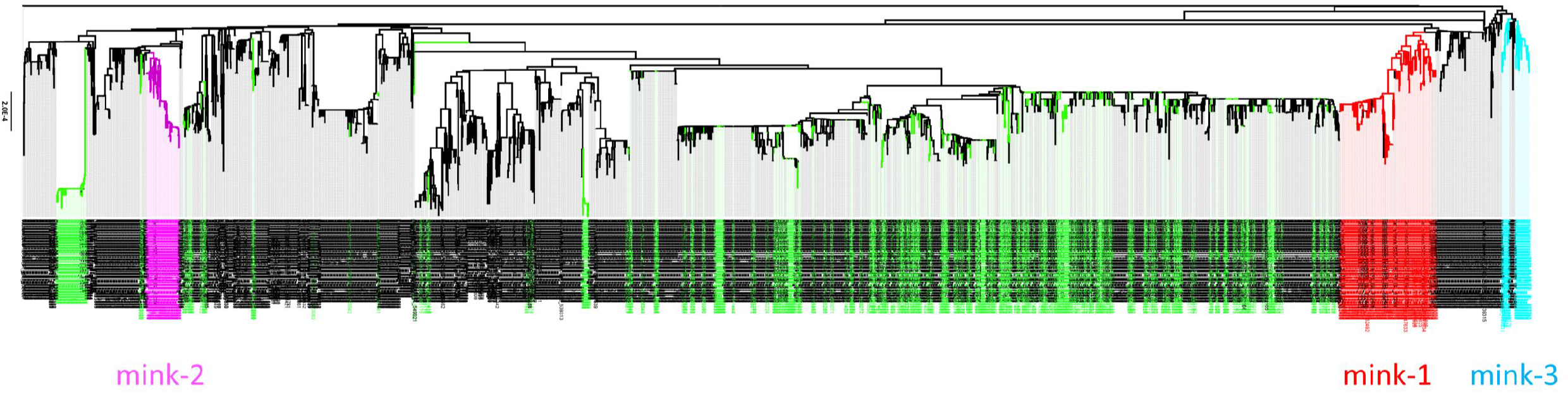
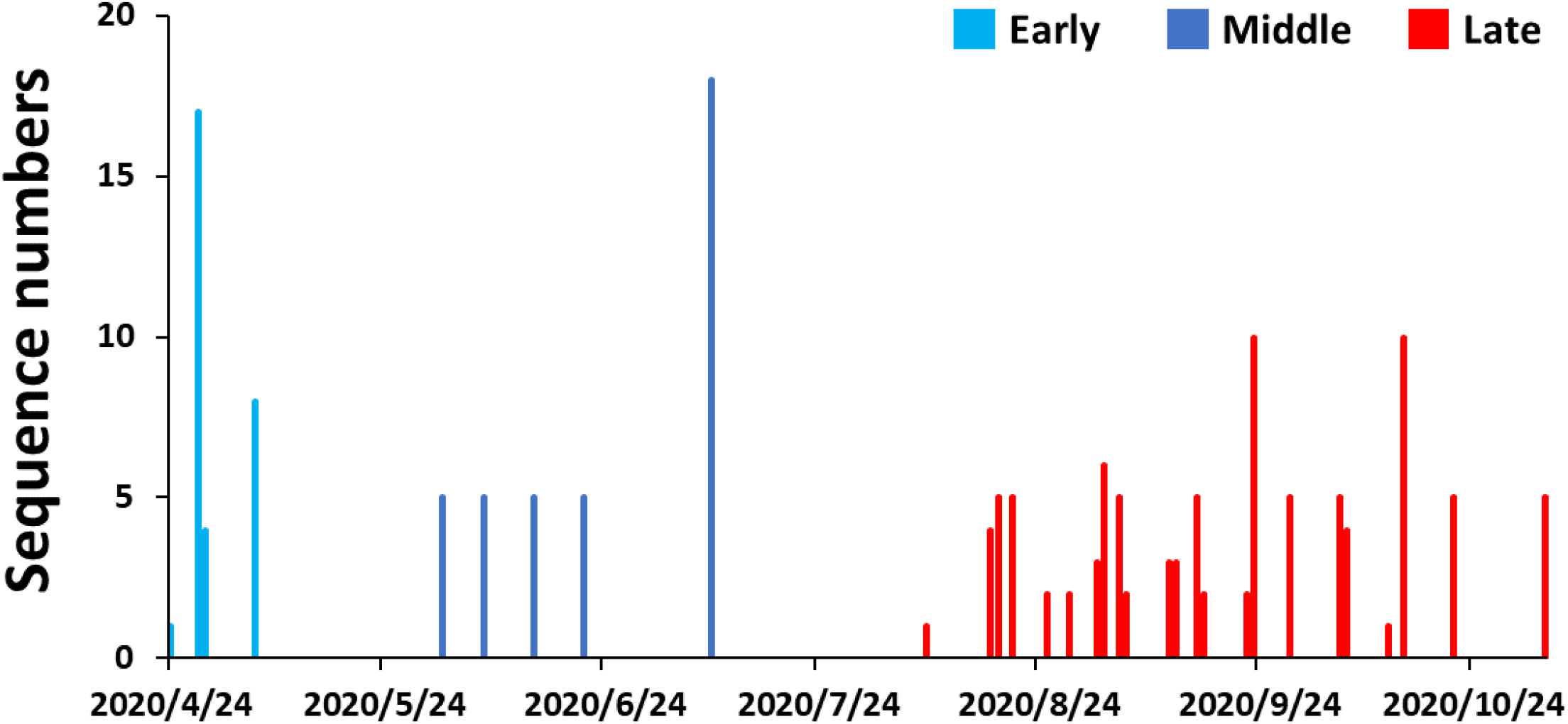
Phylogeny and epidemiology of SARS-CoV-2 that infected minks and humans. (a) The maximum likelihood phylogeny of SARS-CoV-2 derived from minks (colored sequences) and associated humans as of 2020/12/31. (b) Distribution of case numbers in mink-1.

Consistent with the above scenario, strong evidence of positive selection was found in the mink-1 clade as Ka/Ks at the spike protein locus is 5.33 (Table 1, Table S1). Several previous studies have demonstrated that data collected over short time scales may yield biased estimates for Ka/Ks [30–35]. This is either because sufficient time has not passed for natural selection to purge slightly deleterious mutations from the gene pool or because not enough mutations have accumulated to correctly compute a Ka/Ks ratio. Consequently, Ka/Ks ratio should not be used as the sole evidence of positive selection during this short period of time. Nevertheless, not only a higher number of nonsynonymous than synonymous mutations was found in the spike protein, but these mutations were also concentrated in the domains critical for infection. Four of the seven amino acid changes within the spike protein are in the receptor binding domain (RBD) (p = 0.013; Fisher exact test), and three of these four mutations are in the receptor binding motif (RBM) (p = 0.004) (Table 2, Table S2).

**Table 1.** Ka, Ks, and Ka/Ks of different open reading frames of SARS-CoV-2 in different lineages

|  | *All* |  | *orf1a* |  | *orf1b* |  | *S* |  |
| --- | --- | --- | --- | --- | --- | --- | --- | --- |
|  | *Ka x 10<sup>-4</sup>* | *Ks x 10<sup>-4</sup>* | *Ka x 10<sup>-4</sup>* | *Ks x 10<sup>-4</sup>* | *Ka x 10<sup>-4</sup>* | *Ks x 10<sup>-4</sup>* | *Ka x 10<sup>-4</sup>* | *Ks x 10<sup>-4</sup>* |
|  | *Ka/Ks* |  | *Ka/Ks* |  | *Ka/Ks* |  | *Ka/Ks* |  |
| **mink-1 (n=163)** | 2.71 | 5.00 | 2.27 | 2.19 | 0.97 | 4.84 | 5.75 | 1.08 |
|  | 0.54 |  | 1.04 |  | 0.20 |  | **5.33\*** |  |
| **mink-1 Early Phase (n=30)** | 1.86 | 2.49 | 2.39 | 1.00 | 0.10 | 2.35 | 2.55 | 0.87 |
|  | 0.75 |  | **2.39\*** |  | 0.04 |  | **2.93\*** |  |
| **mink-1 Middle Phase (n=38)** | 2.33 | 3.79 | 2.07 | 3.57 | 1.07 | 2.76 | 2.44 | 0 |
|  | 0.61 |  | 0.58 |  | 0.39 |  | 0.64 |  |
| **mink-1 Late Phase (n=95)** | 1.11 | 1.95 | 1.31 | 1.83 | 0.17 | 3.18 | 0.35 | 1.57 |
|  | 0.57 |  | 0.72 |  | 0.05 |  | 0.22 |  |
| **mink-2 (n=59)** | 2.14 | 4.69 | 0.93 | 3.64 | 0.51 | 4.59 | 3.71 | 1.57 |
|  | 0.46 |  | 0.26 |  | 0.11 |  | **2.37\*** |  |
| **mink-3 (n=41)** | 1.92 | 4.29 | 1.63 | 3.01 | 1.32 | 2.74 | 2.86 | 5.66 |
|  | 0.45 |  | 0.54 |  | 0.48 |  | 0.51 |  |
| **B.1.351 (n=783)** | 2.64 | 5.90 | 2.37 | 5.97 | 1.31 | 6.56 | 5.18 | 3.64 |
|  | 0.45 |  | 0.40 |  | 0.20 |  | **1.42\*** |  |
| **B.1.351 with 215G in Spike (n=666)** | 2.25 | 5.44 | 1.79 | 5.91 | 1.38 | 5.55 | 3.63 | 3.22 |
|  | 0.41 |  | 0.30 |  | 0.25 |  | 1.13 |  |
| **B.1.1.7 (n=1824)** | 1.30 | 2.27 | 1.30 | 2.55 | 1.25 | 1.81 | 0.85 | 1.77 |
|  | 0.57 |  | 0.51 |  | 0.69 |  | 0.48 |  |
| **Early Stage Human (n=1476)** | 1.76 | 4.07 | 1.35 | 4.66 | 1.17 | 3.55 | 1.93 | 2.13 |
|  | 0.43 |  | 0.29 |  | 0.33 |  | 0.91 |  |

Early stage SARS-CoV-2 are sequences from 12/2019 – 2/2020
\* p < 0.05.

**Table 2.** The distribution of mutation within spike in different lineages of SARS-CoV-2

|  | Outside RBD<br>(1090 a.a) | Outside<br>RBM<br>(1216<br>a.a) | Within RBD<br>(126 a.a) | Within RBM<br>(69 a.a) | Fisher exact<br>p-value<br>(RBD / RBM) |
| --- | --- | --- | --- | --- | --- |
| **mink-1** | 3 | 4 | V367F | Y453F, F486L, N501T | **0.01 / < 0.01** |
| **mink-1 early phase** | 0 | 1 | V367F | F486L, N501T | **< 0.01 / < 0.01** |
| **mink-1 middle phase** | 2 | 2 | - | Y453F | 0.39 / 0.15 |
| **mink-1 late phase** | 2 | 2 | - | F486L | 0.39 / 0.15 |
| **mink-2** | 1 | 1 | - | L452M, F486L | 0.06 / **< 0.01** |
| **mink-3** | 1 | 1 | - | Y453F, F486L | 0.06 / **< 0.01** |
| **B.1.351** | 38 | 40 | V382L, P384L |  | 0.10 / 0.27 |
| **B.1.1.7** | 15 | 15 |  | A475V, S494P | 1.00 / 0.24 |
| **Early stage SARS-CoV-2** | 29 | 30 | V367F | V483A | 0.30/ 1.00 |

RBD: Receptor binding domain
RBM: Receptor binding motif
The RBM is included within RBD. Thus, the mutations listed within RBM are also in RBD.

To further search for evidence of positive selection, we used the fixed effects likelihood (FEL) [28] and adaptive branch-site random effects likelihood mixed effects models (aBSREL) [29] implemented in HyPhy [27]. We only used non-redundant high-quality sequences for these analyses (Fig. S1) (Materials and Methods). The FEL method identified eight codons putatively under positive selection (p < 0.05) within the mink-1 clade (Fig. 2a). Based on epidemiological data, the course of this outbreak in minks was divided into early (April to May), middle (June and July), and late (August to November) phases (Fig. 1b). The positively selected lineage identified by the aBSREL method is the lineage leading to middle and late phases (Fig. 2b). In addition, seven putatively selected sites gradually increased in frequency from early and were finally fixed in the late phase (Fig. 2b). The above observations demonstrate that SARS-CoV-2 gradually acquired adaptive changes to effectively transmit among minks.

**Fig. 2.**
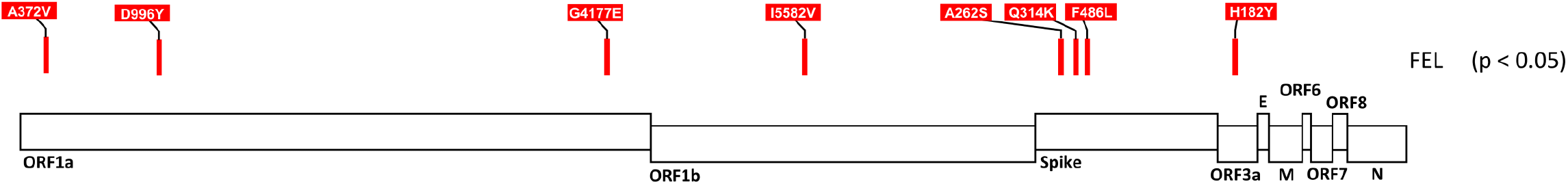
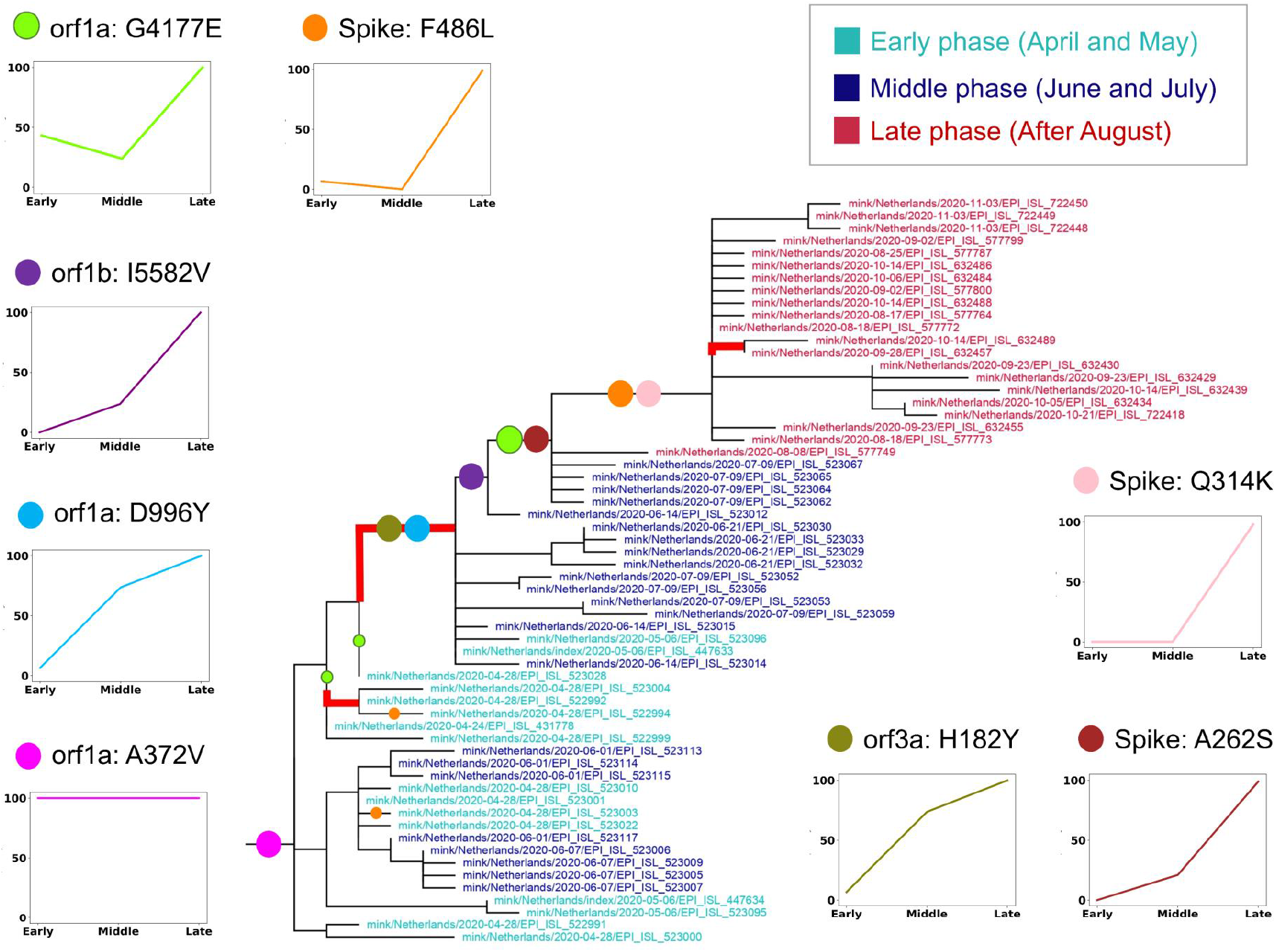
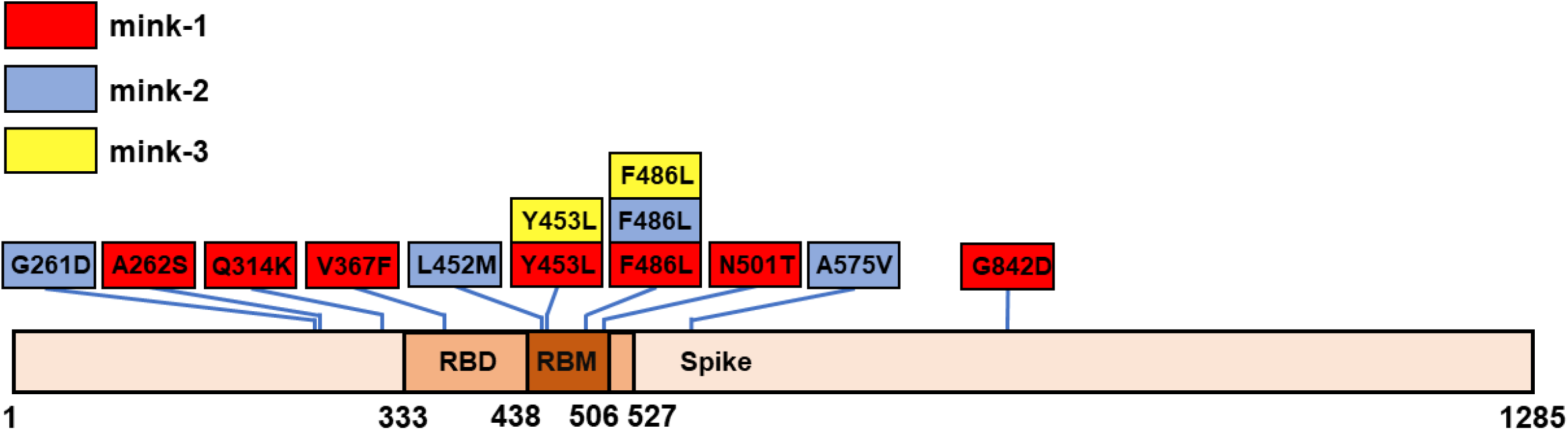
Signature of positive selection in the mink-1 lineage. (a) Amino acid sites putatively under positive selection identified by the fixed effects likelihood (FEL) method implemented in HyPhy. (b) Phylogeny of sequences from the mink-1 lineage. Only non-redundant sequences are included. Red branches are the positively selected lineage identified by the adaptive branch-site random effects likelihood method implemented in HyPhy. Sites putatively under positive selection identified by the FEL method are indicated along the branches with their frequencies in different phases shown. (c) Mutations within the spike protein across clusters of minks.

When Ka/Ks was calculated separately, the whole genome Ka/Ks was highest in the early phase (0.75) and gradually decreased to 0.61 and 0.57 at middle and late phases, respectively (Table 1). Focusing on individual genes, we find that only orf1a and spike protein in the early phase have Ka/Ks > 1. Some short open-reading frames (ORFs) occasionally have Ka/Ks > 1 but that is because their Ks = 0, thus the evidence of positive selection on these ORFs is in doubt (Table S1). Within the spike protein, three amino acid mutations in the early phase are all in the RBD (p < 0.01) and two are in the RBM (p < 0.01) (Table 2, Table S3). This concentration of mutations within RBD/RBM was not seen in the middle or late phases.

In addition to mink-1, signatures of positive selection were also found two other clusters, Mink-2 and −3. The Ka/Ks of the spike protein in mink-2 was 2.37 in Mink-2 and 0.51 in Mink-3 (Table 1 and Table S1). Consistently, both mink-2 and −3 lineages have mutations concentrated in the RBM of the spike protein (p < 0.01; Table 2 and Table S4). It is noteworthy that two mutations within the RBM, Y453F and F486L, probably optimize spike binding affinity to the mink ACE2-receptor (Fig. 2c) [36] and repeatedly occurred on different mink lineages. The convergence of identical mutations from different lineages provides strong evidence of positive selection and implies the adaptation of the virus to its new mink hosts.

### Hitchhiking Under Positive Selection

We found a strong signature of positive selection on the mink-1 lineage. When different phases of the epidemic were analyzed separately, evidence of adaptive evolution was most prominent during the early phase, but scarce in the middle and late phases (Fig. 2b, Table 1).

Nevertheless, the effect of positive selection can leave a trace on linked neutral variation, i.e., the linked variation hitchhikes to either low or high frequencies. Although the frequency distribution of variation can be influenced by several evolutionary processes, an excess of derived variants at high frequency is a unique pattern produced by genetic hitchhiking due to positive selection [37]. We thus constructed site frequency spectra (SFSs) of both synonymous and nonsynonymous changes to look for evidence of positive selection.

SFSs of both synonymous and nonsynonymous changes in mink-1 were skewed toward high frequency mutations (Fig. 3a). Assuming the SARS-CoV-2 population has grown exponentially [16, 19], the observed SFSs significantly deviated from neutral expectation under this population growth model [25] (Materials and Methods), further supporting the idea that the SARS-CoV-2 that transferred from human to mink hosts experienced strong positive selection.

**Fig. 3.**
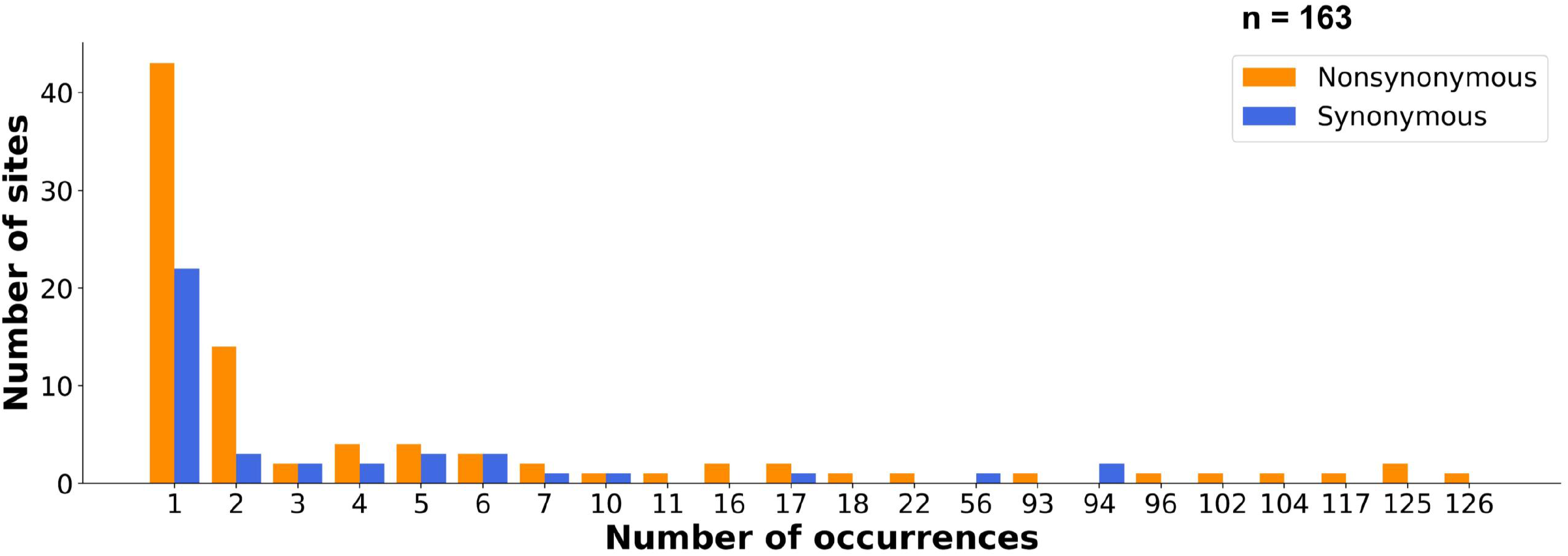
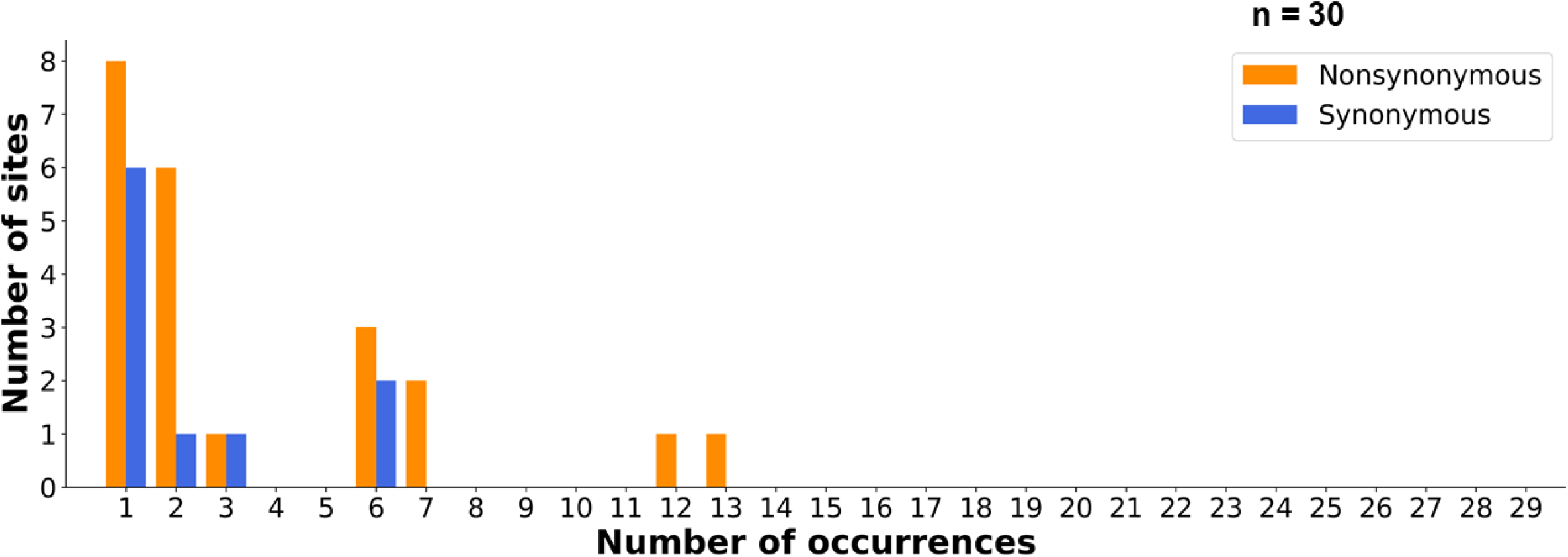
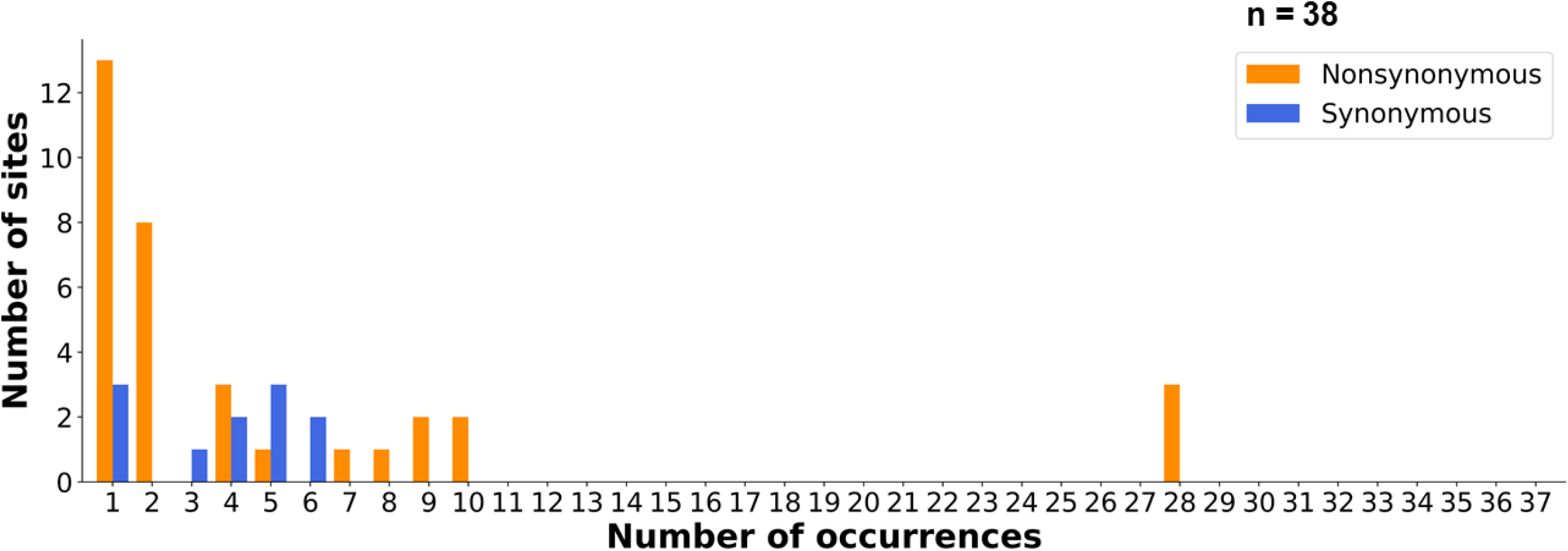
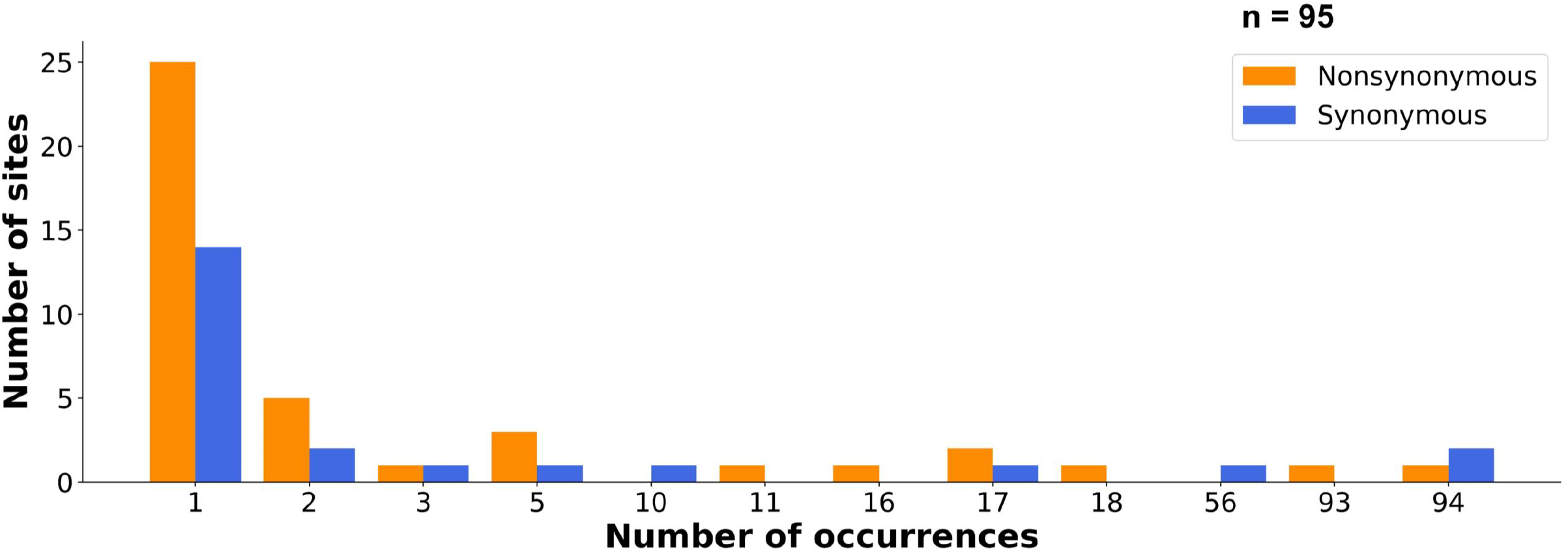
Site frequency spectra of SARS-CoV-2 in the mink-1 lineage in each epidemic phase. (a) Site frequency spectra (SFSs) of the mink-1 lineage. Significant deviation from neutral expectation under exponential population growth was found in both synonymous (p < 10^-3^) and nonsynonymous mutations (p < 10^-5^). (b) SFSs of the mink-1 lineage in the early phase. Neither synonymous nor nonsynonymous mutations deviate from the neutral expectation. (c) SFSs of mink-1 in middle phase. Only nonsynonymous mutations are deviated from neutral expectation (p < 10^-4^). (d) SFSs of the mink-1 lineage in the late phase. Significant deviation from neutral expectation is seen in both synonymous (p < 10^-5^) and nonsynonymous (p < 10^-3^) mutations.

Analyzing each phase separately, we find no excess of high frequency mutations at either nonsynonymous (p = 0.23) or synonymous sites (p = 0.27) (Fig. 3b) in the early phase. That is because the power to detect genetic hitchhiking is compromised when the frequency of advantageous mutations is low [38–40]. As shown in Fig. 2b, putative sites under selection were in low frequency, resulting in a lack of detectable deviation from neutrality.

When advantageous mutations reach high frequency in a population, the SFS reflect a deviation from neutral expectation [38]. Consequently, it is reasonable to expect the excess of high frequency mutations in the middle and late phases of the outbreak as shown in Fig. 3c and d. If we further divide the late phase into late phase I (August and September) and late phase II (October and November), the hitchhiking effect is most prominent in late phase I (Fig. S2a) but less so in late phase 2 (Fig. S2b), demonstrating a rapid decay in the signature of positive selection.

### Evolution of SARS-CoV-2 during the early epidemic

Weak signs of adaptive evolution during the early phase of the epidemic of SARS-COV-2 in humans have been observed in many studies [15, 16, 19, 41]. The Ka/Ks in the whole genome and the spike protein before 2/29/2020 are 0.43 and 0.91 (Table 1), respectively. In addition, very few mutations occurred in the RBD or RBM of the spike protein (Table 2, Table S5). It is possible that the virus may have experienced a recent episode of adaptive evolution before the outbreak. For example, during the short episode of SARS-CoV outbreak in 2002-2003, the Ka/Ks ratio of its spike gene was highest (1.248) in the early phase, but reduced to 0.219 during the late phase [8]. Nevertheless, the SFS still showed signs of genetic hitchhiking due to positive selection (Fig. S3), similar to what we found in the late stage of mink-1.

As shown in Fig. 4a, SFSs of both synonymous and nonsynonymous changes were skewed toward high frequency, which may suggest a signature of positive selection. However, the pattern should be interpreted with caution. The results shown in the Fig 4a were based on an outgroup comparison. The divergence at synonymous sites between SARS-CoV-2 and RaTG13 was 17%, approximately three-fold greater than between humans and rhesus macaques [42]. Indeed, it has been illustrated that inferring directionality of changes via the bat outgroup does not appear to be credible [43]. With such high level of divergence, the possibility of multiple substitutions cannot be ignored [44], especially since substitutions in coronavirus genomes are strongly biased toward transitions [45, 46].

**Fig. 4.**
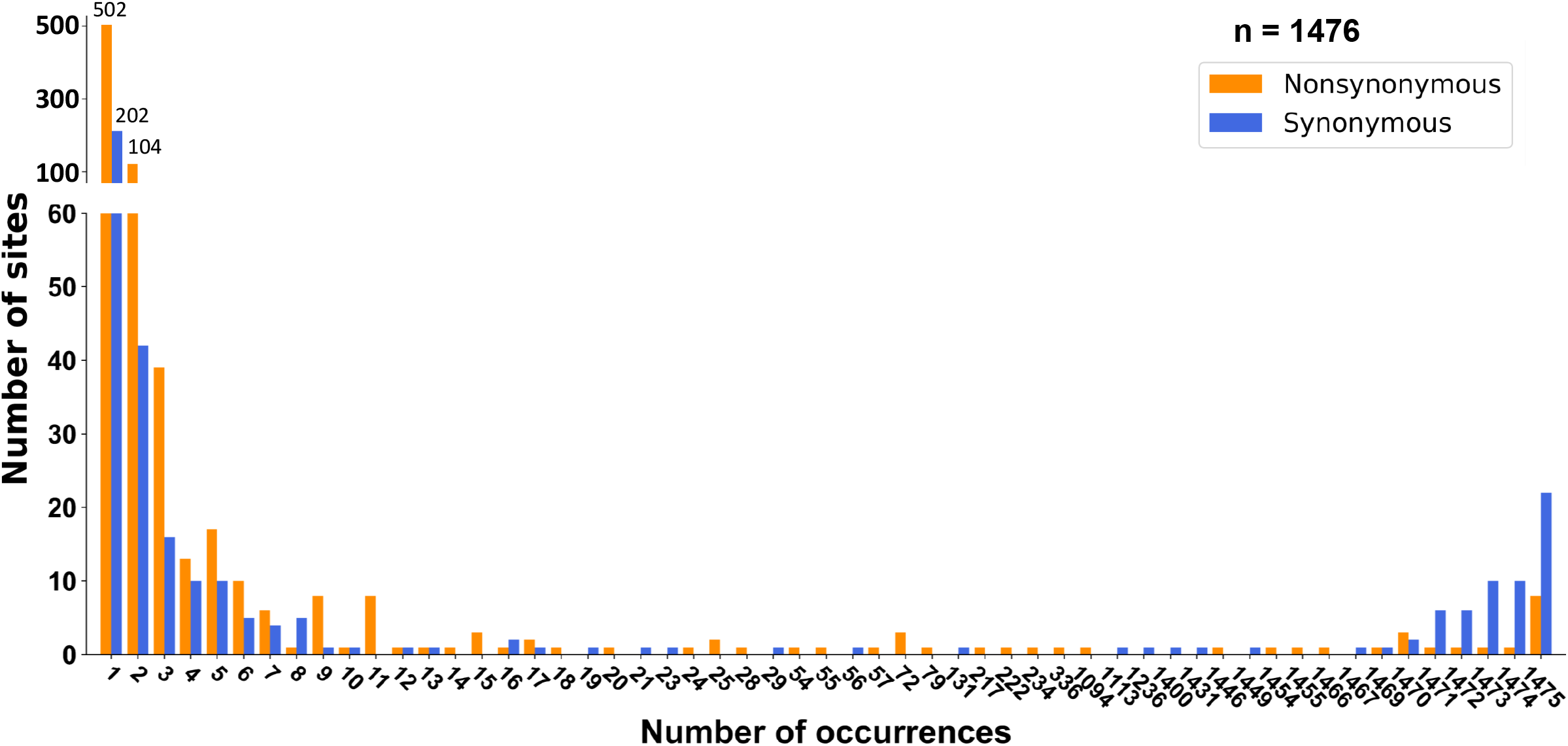
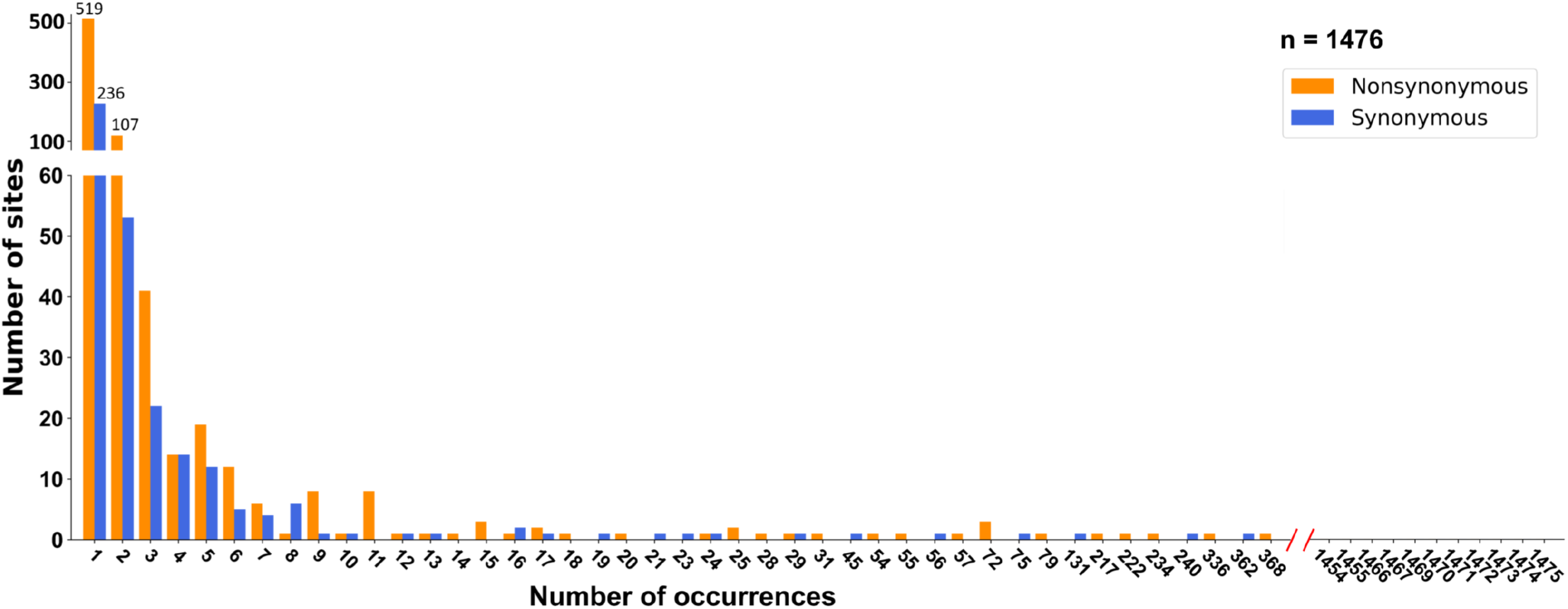
Site frequency spectra of SARS-CoV-2 during the early epidemic (2019/12-2020/2) in humans. (a) Site frequency spectra (SFSs) inferred using RaTG13 as the outgroup. Significant deviation from neutral expectation under exponential population growth was found in both synonymous (p < 10^-5^) and nonsynonymous mutations (p < 10^-5^). (b) SFSs cross-referenced by the phylogeny and date of sampling (see main text for details). Neither synonymous nor nonsynonymous mutations deviate from the neutral expectation.

Among all mutations in Fig. 4a, 32.88% of the changes were C to T transitions (Table 3a), two-fold higher than T to C transitions (15.02%). The bias toward C to T changes was probably mediated either through selective pressures by a CpG-targeting mechanism involving the Zinc finger Antiviral Protein (ZAP), C to U hypermutation by APOBEC3 cytidine deaminases, or escape from the host immune system [47–52].

**Table 3.** Directionality of nucleotide changes of SARS-CoV-2 in (a) polymorphism (early stage (12/2019 – 2/2020)) and (b) divergence

| (a) | Outgroup comparison |  |  | Adjusted |  |
| --- | --- | --- | --- | --- | --- |
|  | Changes | # | (%) | # | (%) |
|  | C -> T | 381 | 32.88% | 420 | 36.24% |
|  | T -> C | 174 | 15.02% | 132 | 11.39% |
|  | A -> G | 147 | 12.69% | 134 | 11.57% |
|  | G -> T | 144 | 12.43% | 149 | 12.86% |
|  | G -> A | 99 | 8.55% | 107 | 9.24% |
|  | A -> T | 46 | 3.97% | 43 | 3.72% |
|  | T -> A | 44 | 3.8% | 49 | 4.23% |
|  | C -> A | 32 | 2.77% | 32 | 2.77% |
|  | A -> C | 31 | 2.68% | 30 | 2.59% |
|  | T -> G | 29 | 2.51% | 29 | 2.51% |
|  | G -> C | 18 | 1.56% | 20 | 1.73% |
|  | C -> G | 14 | 1.21% | 14 | 1.21% |

| (b) | Ancestor to SARS2 |  |  | Ancestor to RaTG13 |  |
| --- | --- | --- | --- | --- | --- |
|  | Changes | # | (%) | # | (%) |
|  | T->C | 185 | 38.07% | 212 | 37.53% |
|  | C->T | 112 | 23.05% | 137 | 24.25% |
|  | A->G | 75 | 15.44% | 83 | 14.7% |
|  | G->A | 40 | 8.24% | 51 | 9.03% |
|  | T->A | 26 | 5.35% | 26 | 4.61% |
|  | A->T | 14 | 2.89% | 25 | 4.43% |
|  | C->A | 10 | 2.06% | 8 | 1.42% |
|  | T->G | 7 | 1.45% | 8 | 1.42% |
|  | A->C | 7 | 1.45% | 7 | 1.24% |
|  | G->T | 6 | 1.24% | 4 | 0.71% |
|  | C->G | 3 | 0.62% | 3 | 0.54% |
|  | G->C | 1 | 0.21% | 1 | 0.18% |

Strikingly, the changes from the common ancestor to SARS-CoV-2 and RaTG-13 were strongly biased toward T to C, 50% higher than C to T (Table 3b). Contrasting patterns between divergence and polymorphism imply that many nucleotide sites may have been changed back and forth during evolution, further increasing the complexity of inferring directionality of change using outgroups. For example, the earliest available SARS-CoV-2 genome (2019/12/24) has the characteristic motif C:C:T at the nucleotide residues 8 782, 18 060, and 28 144, different from the RaTG13 **T:T:C** sequence (Table S6). The first mutant strain carrying the **T:**C:**C** motif, with the bold SNVs matching RaTG13, was found on 2019/12/30. At the same time another 20 genome sequences carrying the C:C:T motif were recovered in Wuhan, China. The first viral strain carrying the **T:T:C** motif was found in the USA on 2020/1/19 and Guangdong 2020/1/20. Therefore, the **T:T:C** motif, although matching RaTG13, is composed of derived instead of ancestral nucleotides.

To get round the potential problem of multiple mutations, we cross-referenced the phylogeny and date of sampling [15]. Many T to C changes based on outgroup comparison were inferred as C to T changes (Table 3a), and all mutations listed as high frequency in Fig. 4a were re-assigned to the other side of the frequency spectra (Fig. 4b). The re-estimated SFSs now only show an excess of low frequency mutations, consistent with a recent origin of SARS-CoV-2 and suggesting that population expansion plays the main role during the evolution of this virus. We did not observe a significant deviation from neutral expectation of growing populations. Consequently, as shown in many studies, SARS-CoV-2 had mainly been evolving under constraint during the early pandemic [15, 19].

### Test for positive selection within humans

Contrasting patterns between early human sequences and three mink lineages implies they are at different stages of host adaptation. However, the lack positive selection signatures in early human samples does not indicate an absence of adaptive evolution of SARS-CoV-2 in humans. Many studies show evidence of adaptation of the spike protein and other genes, especially in the lineages emerging during the last quarter of 2020, including lineage B.1.1.7 (α strain) and B.1.351 (β strain) [16, 17, 53, 54].

On the branch leading to the B.1.351 lineage (Fig. 5a), all 12 lineage defining codons are putatively under positive selection according to the FEL method (p ≤ 0.002). Additionally, two out of five amino acid changes in the spike protein are within the RBM (p =0.05). While the B.1.351 was identified in October 2020 [54], many of lineage defining mutations accumulated step by step during August and September of 2020 (Fig. 5a). Consequently, like mink-1, the B.1.351 lineage also gradually acquired adaptive changes during its evolution.

**Fig. 5.**
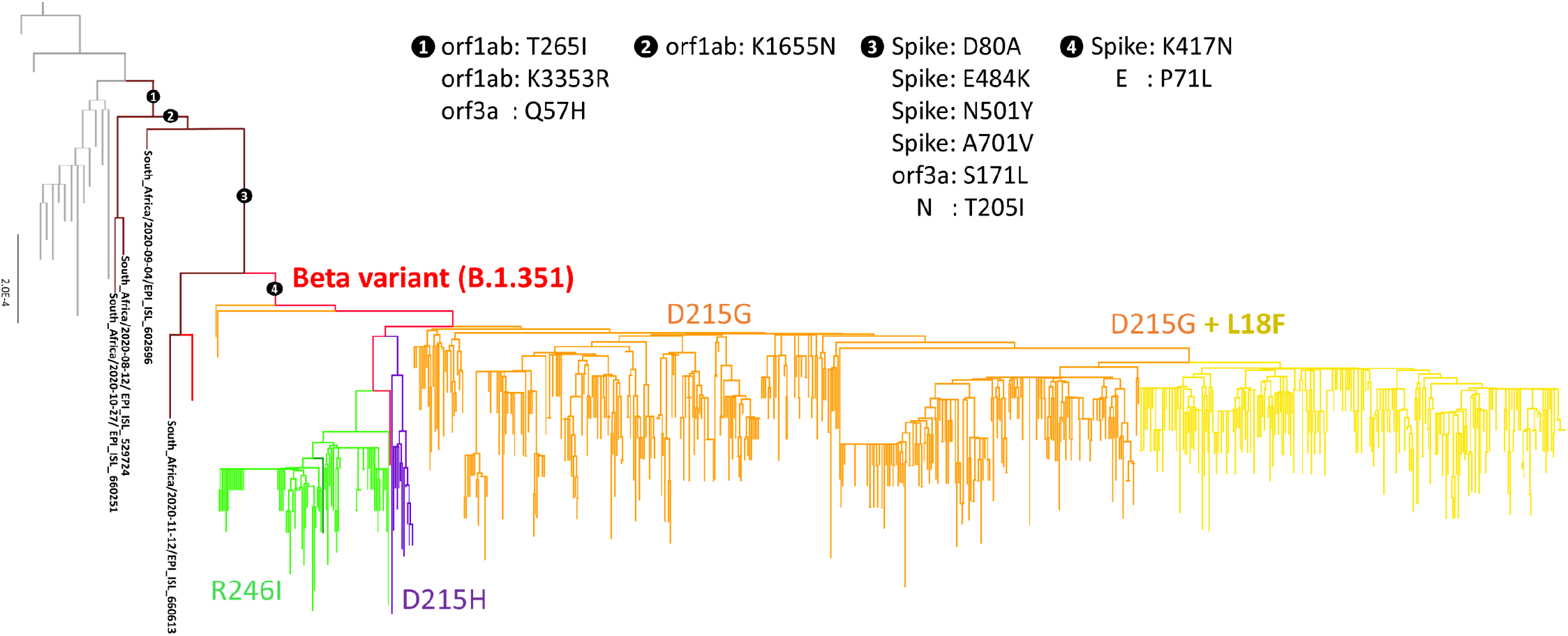
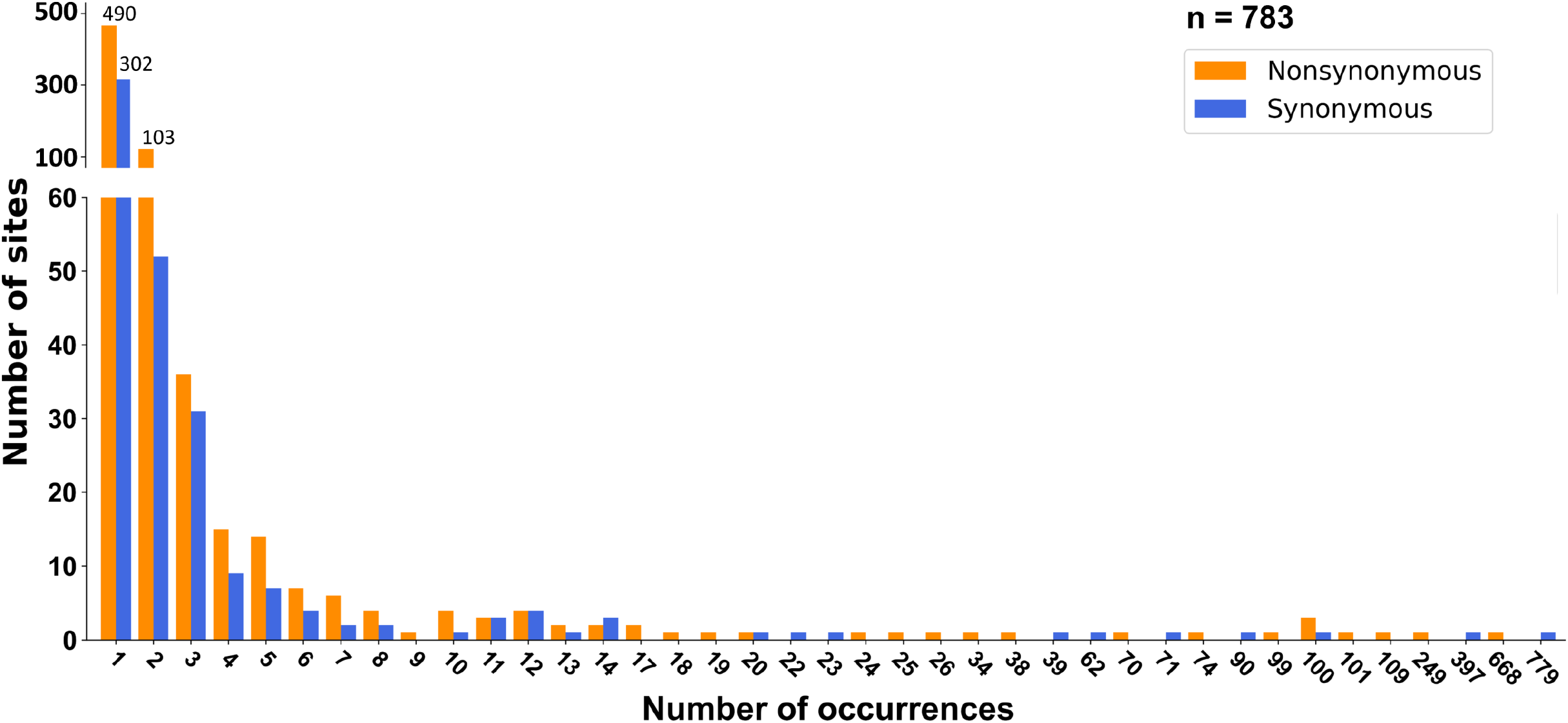
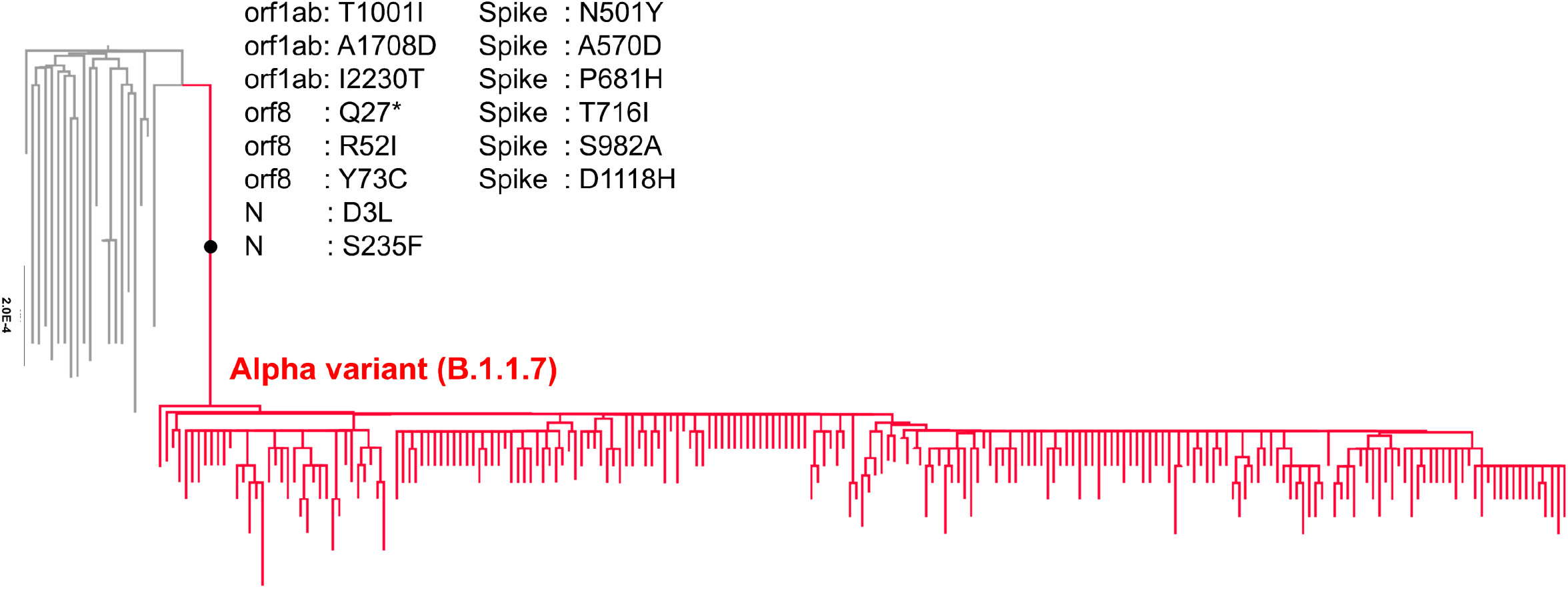
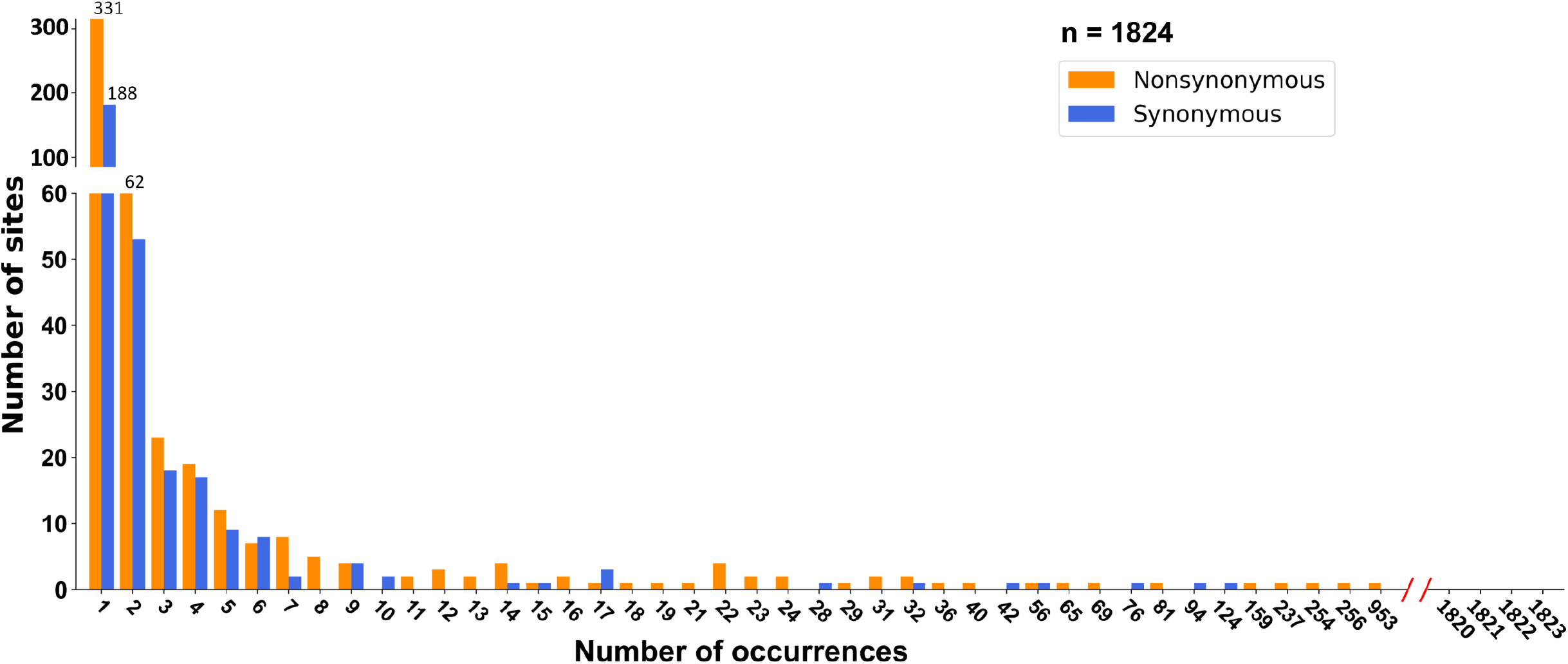
Signatures of positive selection in B.1.351 (β) and B.1.1.7 (α) strains of SARS-CoV-2 (a) Phylogeny of B.1.351. The branches tested for positive selection by the fixed effects likelihood (FEL) method are labeled with numbers and putative selected amino acid sites on the specific branches are listed. (b) Site frequency spectra (SFSs) of B.1.351. Both nonsynonymous (p = 0.03) and synonymous (p < 10^-2^) mutations deviate from the neutral expectation. (c) Phylogeny of B.1.1.7. The branches tested for positive selection by fixed effects likelihood (FEL) method are labeled with a dot and putative selected amino acid sites on the specific branches are listed. (d) Site frequency spectra (SFSs) of B.1.1.7. No evidence of genetic hitchhiking was found in nonsynonymous or synonymous mutations.

Within B.1.351, Ka/Ks of the spike protein was 1.42, partially due to gene specific Ks (3.64 x 10^-4^) much smaller than the genome-wide value (5.90 x 10^-4^). The elevated Ka/Ks ratio was due to co-segregation of D215G and R246I within the spike protein. Although both are considered to be lineage-defining mutations, they are mutually exclusive (Fig. 5a). If we only count sequences carrying 215G, the Ka/Ks of the spike protein is reduced to 1.13, not significantly greater than one (Table 1, p > 0.05).

Nevertheless, positive selection still left a trace on linked neutral variation within B.1.351 (Fig. 5b). The SFSs of B.1.351 show an excess of high frequency synonymous (p = 0.004) and nonsynonymous (p = 0.034) mutations. Significant deviation from neutral expectation in nonsynonymous changes was due to high frequency of the D215G mutation. When only sequences carrying 215G are considered, the SFS skew was only significant for synonymous mutations (p = 0.003), demonstrating the effect of genetic hitchhiking under positive selection (Fig. S4). When each month was analyzed separately, the hitchhiking effect only persisted until November (Fig. S5).

In contrast to lineage B.1.351 and mink-1, both of which show several steps of adaptation with each step including a few amino acid changes, the B.1.1.7 carries many more (13) lineage-specific amino acid changes, all of them putatively under positive selection according to the FEL method (p ≤ 0.01), prior to the detection of this lineage (Fig. 5c). It is hypothesized that the putative adaptation of B.1.1.7 may have resulted from virus evolution within a chronically infected individual [55]. Therefore, this strain seems to be well-adapted at its appearance, a situation similar to the early epidemic of SARS-COV-2 in humans. The Ka/Ks ratios of the whole genome on this lineage is 0.57 (Table 1). In addition, the SFSs did not significantly deviate from neutral expectation of growing populations (Fig. 5d).

## DISCUSSION

### Cryptically circulating SARS-CoV-2 before the 2019 outbreak

Using several approaches, we identified signs of adaptive evolution in the lineages leading to mink-1, B.1.351, and B.1.1.7. Although the signature of positive selection diminished quickly, it still left a trace on linked neutral variation as shown by SFSs in mink-1 and B.1.351. However, as found by many previous studies, we did not detect evidence of positive selection the early episode of the SARS-CoV-2 pandemic [15, 16, 19]. It is possible that SARS-CoV-2 has been cryptically circulating within humans for a sometime before being noticed [56–58]. Alternatively, the virus might have “preadapted” to the human host prior to its emergence [19, 59].

However, it is one thing for a virus to occasionally spill over to a new host, it is another for it to cause an epidemic. For example, although animal influenza viruses occasionally infect humans, many of these zoonoses do not cause subsequent widespread transmission and stop after infecting a few people [60, 61]. In addition, while many studies documented the susceptibility of different animal species to SARS-CoV-2, including cats, dogs, tigers, lions, ferrets, rhesus macaque, and tree shrews [62–68], it only caused outbreaks in minks [69]. Furthermore, as shown in Fig. 1, transmission from humans to minks occurred multiple times, majority of them failing to trigger an epidemic. The above observations indicate that many attempts of SARS-CoV-2 to jump cross species boundaries appeared as transient ‘spillover’ infections and soon died out.

For a virus to adapt to new hosts and to produce a pandemic, it must gain the ability to (1) bind and enter host cells, (2) evade host restriction factors and immune responses, and (3) transmit effectively among hosts. Such adaptation is not likely to have emerged suddenly but, instead, may have evolved step by step with each step favored by natural selection [5–7, 61, 70, 71]. The binding of the spike protein to the host ACE2 receptor is the first and one of the most important steps in SARS-CoV-2 infection of host cells. While the history of early adaptation is unknown, the RBD of the SARS-CoV-2 spike protein appears to be highly specialized to human ACE2 [72]. Substitution of eight SARS-CoV-2 RBD residues proximal to the ACE2-binding surface with those found in RaTG13 is almost universally detrimental to human ACE2 receptor usage [73]. Both SARS-CoV-2 and RaTG13 bind poorly to *R. sinicus* ACE2 [74]. These observations indicate that SARS-CoV-2 is well adapted to humans. We observe in this study a strong signature of positive selection in the viral strains that successfully established continuous infections among minks. By analogy, it appears unlikely that a nonhuman progenitor of SARS-CoV-2 would require little or no novel adaptation to successfully infect humans.

We instead hypothesize that SARS-CoV-2 has been cryptically circulating among humans before the current outbreak. During the period of unawareness, the virus had gradually accumulated adaptive changes that enabled it to effectively infect humans and finally cause the pandemic. After adapting to the new host, most RNA viruses exhibit strong negative selection [20]. As we see in the mink-1 and B.1.351 lineages, it takes four to six months after the emergence of adaptive substitutions for the effect of genetic hitchhiking to dissipate. It is therefore reasonable to expect that the signature of adaptive evolution to have quickly diminished after the early SARS-CoV-2 adaption to humans.

Our hypothesis is supported by the situation found in B.1.1.7. This strain was not tracked until it accumulated 13 amino acid changes putatively under adaptive evolution with no known intermediate, and no sign of positive selection was detected after its emergence. The case of B.1.1.7 clearly demonstrates that the process of virus adaption can be stealthy. Although the exact time the virus jumps to human is difficult to estimate without further information, we may reference the results from minks and SARS-CoV. According to the outbreak in mink-1 and SARS-CoV, the SFSs still deviated from neutral expectation six months after the first case was recorded (Fig. 3 and Fig. S3). Consequently, it is reasonable to conclude that the actual origin of SARS-CoV-2 is at least six months before the first case in Wuhan, China, in early December 2019. We therefore presume that the virus may have associated with humans before June 2019 [56–58]. However, to verify this hypothesis, it is essential to collect archive samples of pneumonia in the Wuhan area for analysis. These data are needed to trace its evolutionary path and reveal critical steps required for effective spreading.

### Why and how the current pandemic occurred

It is not uncommon that the origin of virus infection dates back before the awareness of the epidemic. For example, molecular clock dating suggests the onset of HIV-M and -O epidemics occurred at the beginning of the 20th century [75–77]. However, it was not until 1980 that the virus was finally confirmed as the causal agent of AIDS [78–80]. Because natural selection is working in the original hosts, higher infectiousness of a zoonotic virus in the wild animal hosts than in humans is expected. As viruses evolve they may invade multiple times but fail to trigger an epidemic as shown in the SARS-CoV-2 that infected minks. Occasionally, after many failed attempts, the emergence of a new strain can sustain the infection long enough to acquire new mutations for further enhancement of infectiousness in humans.

Even viruses that appear to be well adapted to humans may fail to induce an outbreak [71]. That is because during the time the virus built up its ability to infect humans, herd immunity can develop in local populations and impede epidemic spread. Such viruses would then be more infectious outside the enzootic area as populations outside of the area are immunologically naïve [5]. That is why the place of origin is not necessary the same as the outbreak location, as can be seen in the cases of HIV and influenza [81–83]. It is also possible that a changing ecological environment may impact virus spread. In the case of the canine influenza virus (CIV), which jumped to dogs in the late 1990s from an equine influenza strain prevalent in horses [84], Dalziel et al., found that CIV is largely confined to dog shelters in the US, where most dogs are infected soon after they arrive. But the virus cannot be maintained for long in smaller facilities or in the companion dog population without input from the larger shelters. These hotspot dynamics give a clear picture of what can happen in the time between the beginning of a host range shift and the onset of a possible pandemic [85].

## ACKNOWLEDGEMENT

We thank Anthony Greenberg for comments. This study was supported by grants from Ministry of Science and Technology (MOST), Taiwan (109-2311-B-002 -023 -MY3, 109-2327-B-002-009, 109-2634-F-002-043) and National Taiwan University, College of Medicine (NSC-131-5).

