## Supplement figures for "Signatures of adaptive evolution during human to mink SARS CoV2 cross-species transmission inform estimates of the COVID19 pandemic timing"

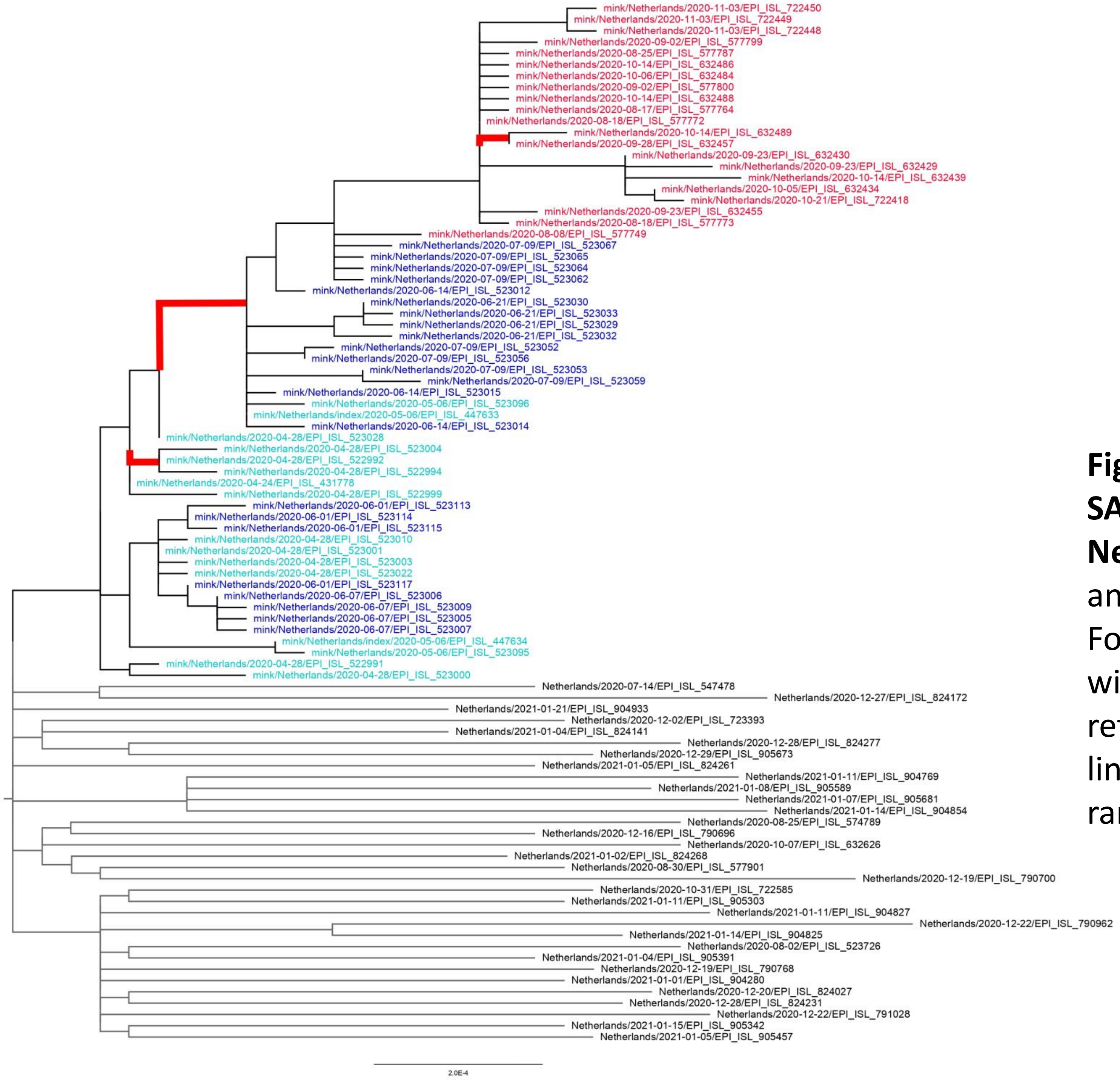


**Fig. S1 The maximum likelihood phylogeny of SARS-CoV-2 of mink-1 and humans from Netherlands.** For mink-1, only non-redundant and no ambiguous site sequences were included. For SARS-CoV-2 from humans only sequences with lesser than 99.9% nucleotide identify were retained. Red branches are positive selected lineage identified by the adaptive branch-site random effects likelihood method.

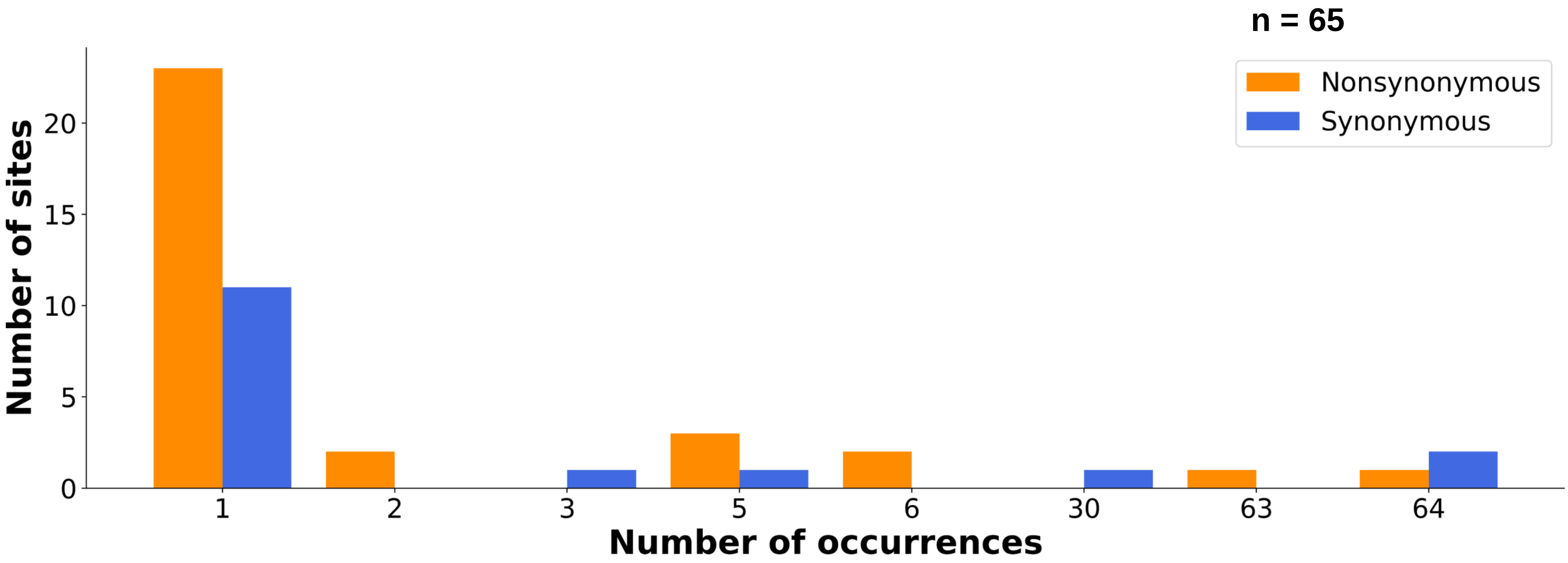


**Fig. S2a. Site frequency spectra of mink-1 during the last phase I of epidemic (August and September 2020) .** Significant deviation from neutral expectation was found in both nonsynonymous and synonymous mutations ($p < 0.01$)

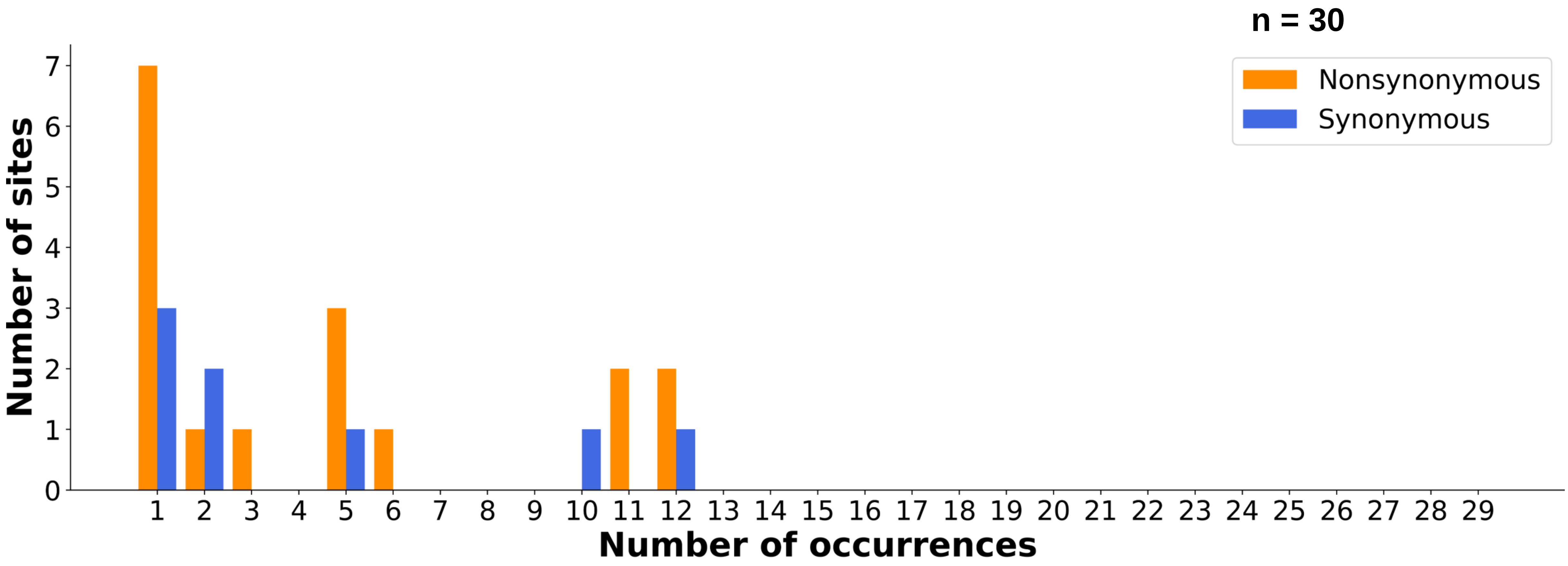


**Fig. S2b. Site frequency spectra of mink-1 during the last phase II of epidemic (October and November 2020) .** Significant deviation from neutral expectation is only found in nonsynonymous mutations (p = 0.02).

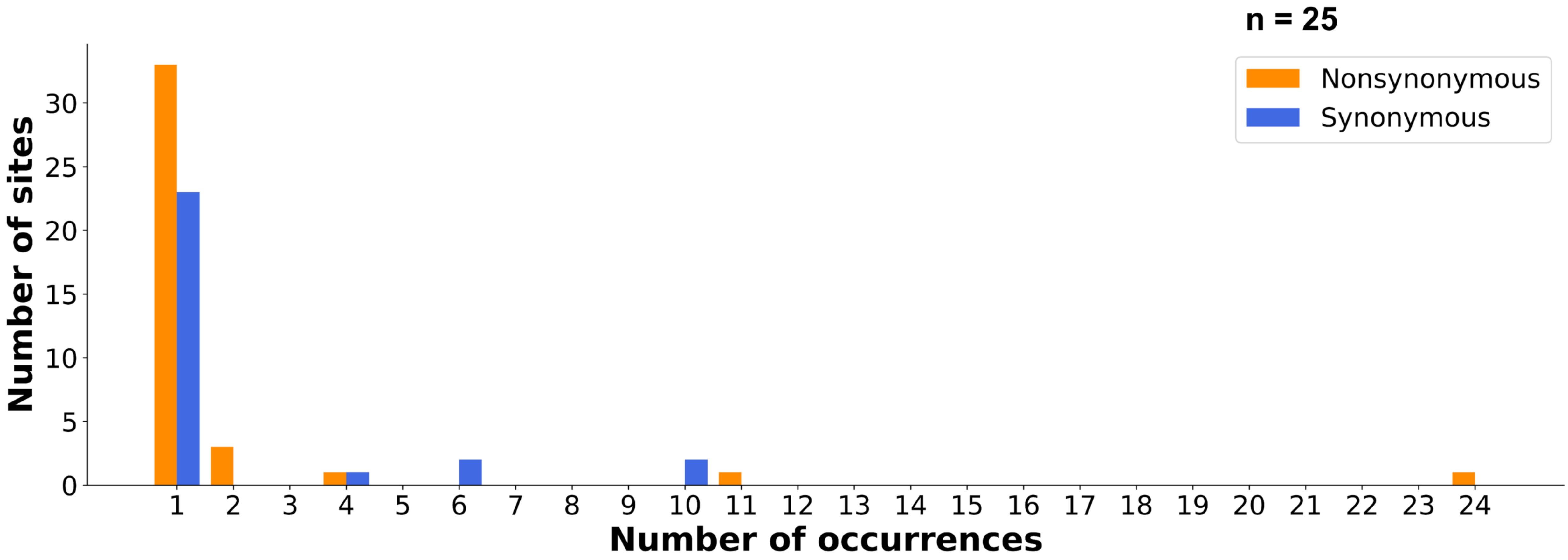


**Fig. S3 Site frequency spectra of SARS-CoV during the last phase of epidemic in 2003.** Significant deviation from neutral expectation was found in nonsynonymous ($p < 0.02$) and synonymous ($p < 0.05$) mutations.

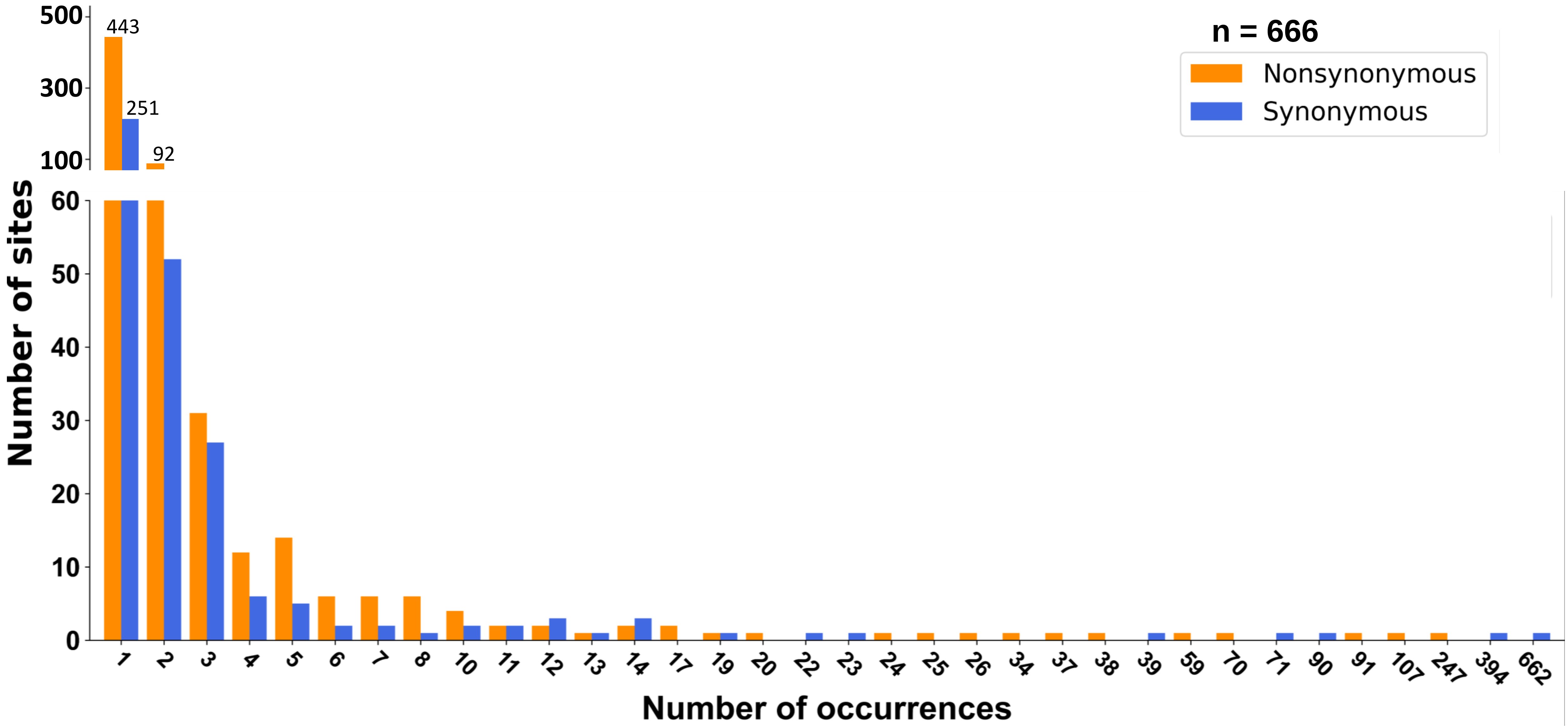


**Fig. S4 Site frequency spectra of B.1.351 with 215G in Spike.** Significant deviation from neutral expectation was only found in synonymous ($p < 10^{-2}$) mutations.

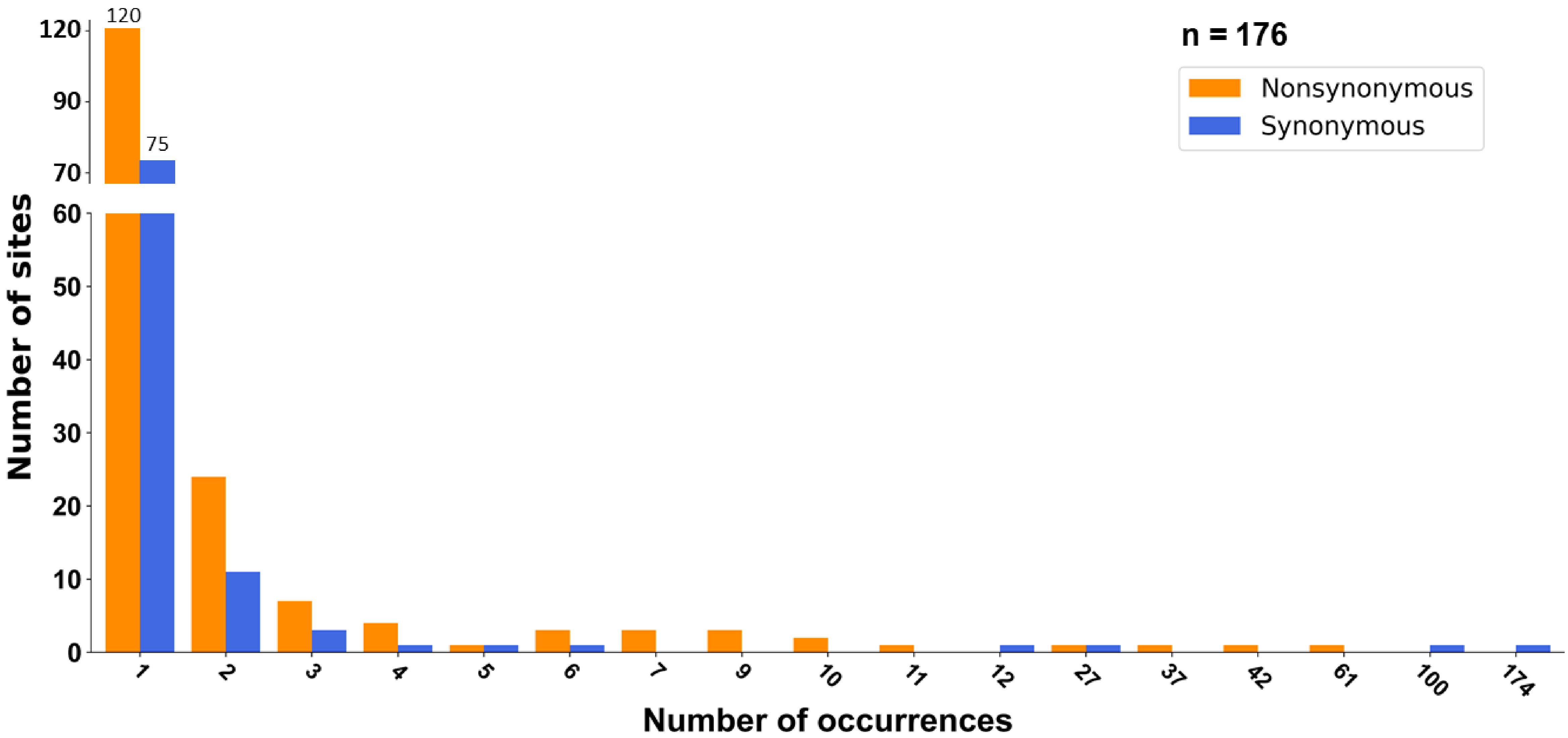


**Fig. S5a Site frequency spectra of B.1.351 strains carrying 215G in Spike in sequences collected in November 2020.** Significant deviation from neutral expectation was only found in synonymous ($p < 10^{-2}$) mutations.

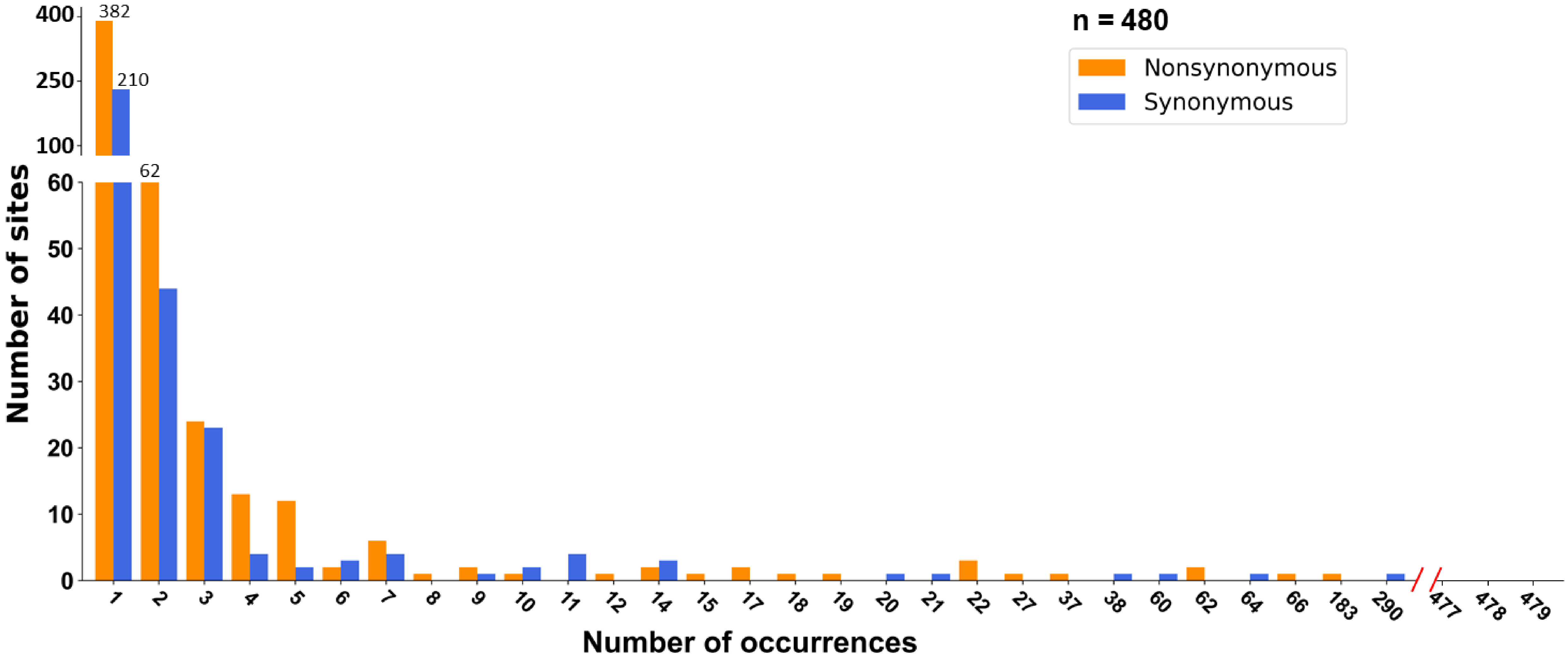


**Fig. S5b Site frequency spectra of B.1.351 strains carrying 215G in Spike in sequences collected in December 2020.** No effect of genetic hitchhiking was.
